# A protective hybrid protein vaccine informed by mapping major histocompatibility class II epitopes in *Cryptococcus neoformans* chitin deacetylases that stimulate CD4^+^ T cell responses

**DOI:** 10.64898/2026.09.07.749995

**Authors:** Charles A. Specht, Lorena V.N. Oliveira, Diana Carlson, Maureen M. Hester, Satya Sahu, Gabriel Kristian Pedersen, Stuart M. Levitz

## Abstract

Infections due to *Cryptococcus* species are estimated to cause over 100,000 deaths a year. No cryptococcal vaccines are approved for human use. We have shown that vaccine formulations consisting of *C. neoformans* chitin deacetylase (Cda)2 and Cda1 adjuvanted with Cationic Adjuvant Formulation 01 (CAF01) protect mice against experimental cryptococcosis by a mechanism dependent upon CD4^+^ T cells. Here, using overlapping peptide libraries, we mapped epitopes in Cda2 and Cda1 that stimulated interferon-gamma (IFNγ) release from splenocytes of C57BL/6 and DR4 mice immunized with these proteins. DR4 mice lack mouse major histocompatibility class II (MHC-II) proteins and express chimeric MHC-II proteins with the specificity of the human HLA-DR allele, HLA-DRB1*04:01. Experimental results were then compared with MHC-II binding predictions using the Immune Epitope Database (IEDB) NetMHCIIpan 4.1 BA MHC-II epitope prediction tool. Unique epitopes in Cda2 and Cda1 were discovered for each mouse strain, some of which were not predicted to bind well to MHC-II. CD4^+^ T cells were responsible for cytokine release as IFNγ production was lost if CD4^+^ T cells were depleted from the splenocyte population. Finally, we tested a hybrid recombinant protein that consisted of portions of Cda2 and Cda1 containing the MHC-II/CD4^+^ T cell epitopes which induce CD4^+^ T cells in DR4 mice. When administered as a CAF01-adjuvanted vaccine, the hybrid protein protected DR4 mice from an otherwise lethal *C. neoformans* pulmonary challenge. These results provide a proof of principle regarding the utility of MHC-II/CD4^+^ T cell epitope mapping in cryptococcal vaccine development.

**Importance:** The development of vaccines to prevent cryptococcosis, a fungal disease which causes over 100,000 deaths annually, is a public health priority. We have demonstrated that adjuvanted vaccines containing two *Cryptococcus neoformans* protein antigens, chitin deacetylase (Cda)2 and Cda1, protect mice against experimental cryptococcosis by a mechanism dependent upon CD4^+^ T cells. In this study, we determined which 15 amino acid peptides of Cda2 and Cda1 stimulate CD4^+^ T cells in C57BL/6 and DR4 mice; the latter mouse strain has human major histocompatibility class II (MHCII) specificity. We then synthesized a small hybrid protein containing those portions of Cda2 and Cda1 that strongly stimulated CD4^+^ T cell responses in DR4 mice. As an adjuvanted vaccine, the hybrid Cda2/1 protein protected DR4 mice from a *C. neoformans* pulmonary challenge. Thus, CD4^+^ T cell/MHCII epitope mapping experimentally validated the design of a protective multi-epitope protein for use in cryptococcal vaccines.

## Introduction

Infections due to encapsulated fungi belonging to the *Cryptococcus neoformans* and *C. gattii* species complexes are estimated to kill over 100,000 people annually (1). The vast majority of cryptococcosis cases occur in people with impaired CD4^+^ T cell function due to factors such as advanced HIV infection, receipt of immunosuppressing medications, and hematological malignancy (2). Most infections are thought to originate following inhalation of airborne fungi; in the absence of effective host defenses, central nervous system dissemination can ensue.

There are no approved cryptococcal vaccines for use in humans. Vaccine formulations consisting of attenuated *C. neoformans* strains and adjuvanted subunit formulations protect mice against experimental cryptococcosis (reviewed in (3–5)). Critical for the development of protective subunit vaccines is antigen and adjuvant discovery. Regarding antigens, we have extensively studied two promising protein candidates, chitin deacetylase (Cda)2 and Cda1. When adjuvanted in glucan particles or with cationic adjuvant formulation 01 (CAF01), Cda2 and Cda1 protect BALB/c, C57BL/6, and DR4 mice from an otherwise lethal *C. neoformans* pulmonary challenge (6–11). The importance of T helper (Th) cells is suggested by data showing protection is lost in mice that lack CD4^+^ T cells or interferon-gamma (IFNγ). Moreover, mice that are vaccinated and infected develop robust antigen-specific IFNγ-producing CD4^+^ T (Th1) cells (6–8, 12).

Development of an adaptive CD4^+^ T cell response requires antigen uptake, processing, and presentation of proteins by antigen-presenting cells (APCs) to the T cell receptor (TCR) (reviewed in (13)). Uptake can occur via phagocytosis, receptor-mediated endocytosis, or pinocytosis. Following proteolytic degradation in endolysosomal compartments, peptides ranging in size from 10-25 amino acids (average length ∼15 amino acids)(14–16) form a complex with major histocompatibility complex Class II (MHC-II) and are transported to the cell surface for presentation. Within the peptide is a 9 amino acid core sequence, of which the first, fourth, sixth and ninth amino acids bind to MHC-II, with the other five amino acids exposed to the TCR on the CD4^+^ T cell (17, 18). The amino acids flanking the core may also contribute to binding (18).

Despite the recognition that adaptive immunity is paramount for host defenses and vaccine-mediated protection against cryptococcosis, information is limited regarding the epitopes in cryptococcal proteins that stimulate antigen-specific CD4^+^ T cell responses. CD4^+^ T cell epitope identification could be useful for development of hybrid multiepitope vaccines and diagnostic tests. Mapping stimulatory MHC-II peptide epitopes within antigenic proteins is challenging in humans because the human leukocyte antigen (HLA) MHC-II is highly polymorphic, and individuals typically have different alleles encoded by the HLA DR, DP and DQ MHC class II loci (13, 19, 20). Immunoinformatic programs can predict the binding affinity of peptides to individual MHC-II molecules (21, 22). While useful, the programs do not account for antigen-processing (such as cleavage by cathepsins and other proteases), posttranslational modifications, T cell precursor frequency, and the T-cell-exposed motifs that bind to the TCR (13, 17, 18, 21, 23). Ultimately, experimental validation of in silico predictions is needed (16).

In the present study, we used overlapping peptide libraries to identify CD4^+^ T cell epitopes present within Cda2 and Cda1 following vaccination of C57BL/6 and DR4 mice. MHC-II peptide presentation in C57BL6 mice occurs via the IAb protein (15). DR4 mice are on the C57BL/6 background, are null for mouse MHC class II, and express the relatively common human HLA-DR allele, HLA-DRB1*04:01 (11, 24). Thus, differences in peptide epitopes that stimulate CD4^+^ T cell responses in C57BL/6 and DR4 mice should be attributable to the disparate MHC-II molecules expressed by the two mouse strains. The experimentally validated epitopes in Cda2 and Cda1 were compared with predicted epitopes using the Immune Epitope Database (IEDB) MHC class II tool, NetMHCIIpan 4.1 BA (22). Finally, a hybrid fusion protein created using regions of Cda2 and Cda1 rich in validated MHC-II/CD4^+^ T cell epitopes protected DR4 mice when used in a vaccine containing a Th1/Th17-inducing adjuvant.

## Results

Use of long peptides to identify MHC-II epitopes in Cda2 and Cda1 stimulating CD4^+^ T <u>cells.</u> In previous experiments, we had synthesized eight “long peptides” (31-35 amino acids in length), which were predicted to have strong binding to the MHC-II H2-IAd allele found in BALB/c mice. When formulated as glucan particle-based vaccines, five of the peptides significantly protected BALB/c mice, but not C57BL/6 mice, from an otherwise lethal pulmonary *C. neoformans* challenge (25). In initial experiments, we immunized DR4 mice with recombinant Cda2 adjuvanted with CAF01. Splenocytes from harvested spleens were prepared and incubated with the eight peptides, and IFNγ was measured in the supernatants (Figure 1A). The nomenclature adopted for the long peptides was PXXL, with P indicating peptide, XX locating the position of the starting amino acid of the peptide in the protein sequence (counting from the N-terminus), and L designating that it is a long peptide. Thus, the 32 amino acid peptide P60L starts at amino acid 60 and ends at amino acid 91. The amino acid sequences of the long Cda2 peptides are shown in Table S1. Of the eight peptides, P60L, clearly stimulated IFNγ above background values. The positive control, recombinant Cda2 protein (Cda2 Prot), was comparably stimulatory.

**Figure 1.**
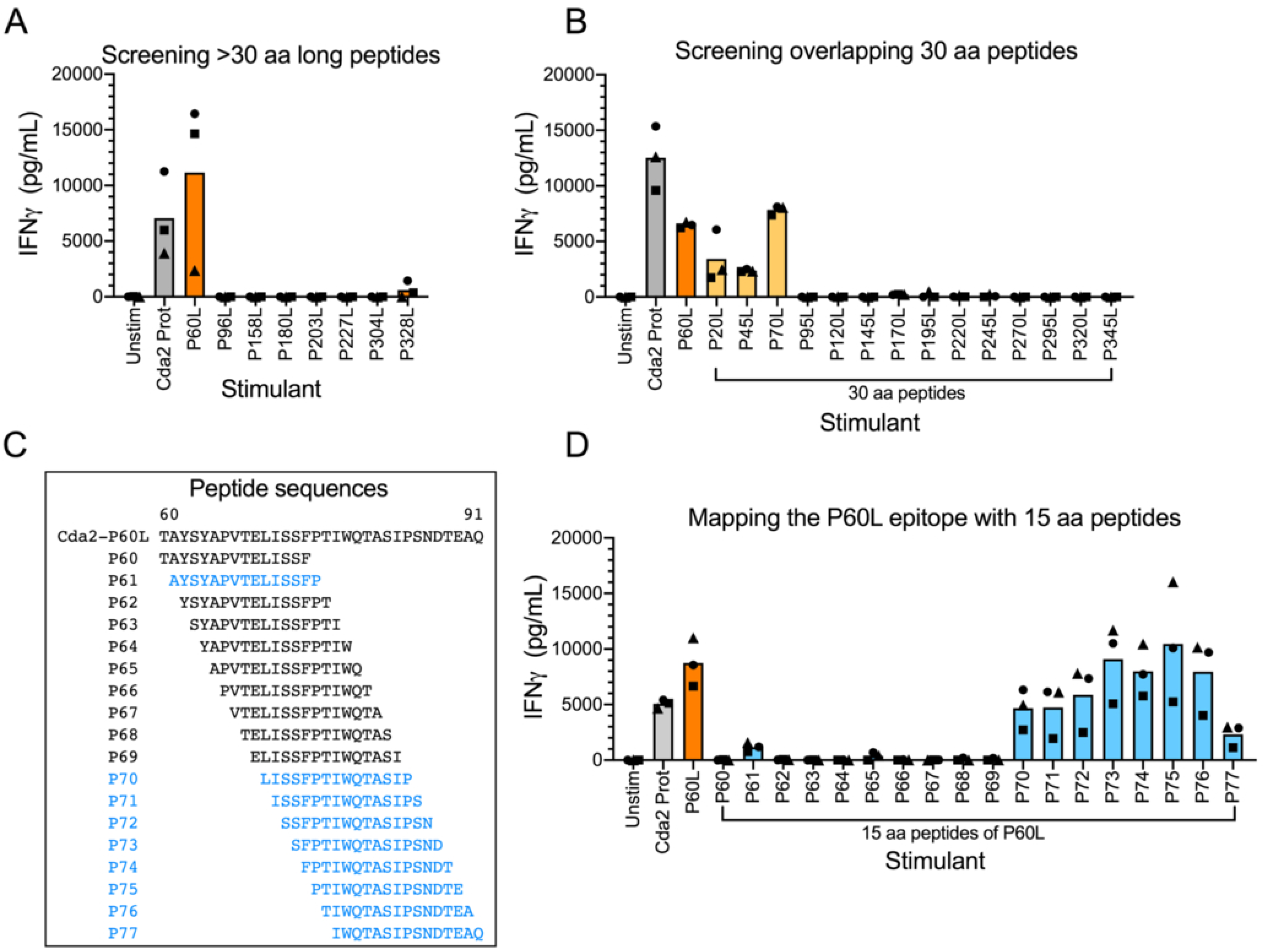
Identification and mapping of a DRB1*0401 epitope of Cda2 using long peptides (≥30 amino acids [aa]). Splenocytes from DR4 mice immunized with recombinant Cda2 protein (Cda2 Prot) were stimulated with the indicated peptides or proteins for three days. IFNγ was then measured in the supernatant. by ELISA. Cda2 Prot was used as the positive control stimulant and unstimulated (Unstim) splenocytes as the negative control. (**A**) Identification of a DRB1*0401 epitope of Cda2 by screening with eight previously published long peptides (25). (**B**) Screening for additional DRB1*0401 epitopes of Cda2 with sequential 30 aa overlapping peptides. (**C**) Sequences of 15 aa peptide used in (**D**) to map the epitope. (**D**) Mapping the location of the epitope identified with Cda2-P60L using 15 aa peptides that span peptide Cda2-P60L. Blue type indicates peptide sequences that were stimulatory while black type denotes peptides that were not stimulatory. Each symbol shape represents results from an individual mouse.

The eight long peptides include the amino acid sequences from only parts of Cda2. We next took an unbiased approach to epitope identification by screening 30 amino acid peptides (30mers) that overlap by five amino acids and span the entire amino acid sequence of recombinant Cda2 (Figure 1B and Table S1). IFNγ release from splenocytes obtained from immunized DR4 mice was stimulated by P20L, P45L, and P70L. The positive control, recombinant Cda2, and the previous hit, P60L, also stimulated IFNγ. To determine which amino acids within the P60L peptide were responsible for stimulating IFNγ production, we synthesized a set of 15 amino acid (15mer) peptides that overlapped by 14 amino acids and encompassed the entire span of P60L (Figure 1C). The nomenclature adopted for the 15mer peptides was PXX, with P indicating peptide; and XX denoting the position of the starting amino acid of the peptide in the protein sequence. Positive hits, as measured by splenocyte IFNγ release, were seen after stimulation of immune splenocytes with peptides P70-P77 (Figure 1D). A very modest response was also seen with P61.

The same approach was taken with C57BL/6 mice immunized with Cda2 (Figure S1A and S1B and Table S1) as had been done with the DR4 mice. IFNγ release was stimulated by P60L and P203L in Figure S1A; and P20L, P45L, P70L, P220L, P295L, P320L, and P345L, as shown in Figure S1B. Lastly, using overlapping 30mers spanning Cda1, we identified CD4^+^ T cell epitopes in DR4 and C57BL/6 mice immunized with recombinant Cda1 (Figure S1C and S1D and Table S2). Two peptides that stimulated IFNγ release, P320L and P345L, were discovered in Cda1-immunized DR4 mice. For the immunized C57BL/6 mice, we also found two peptides, P120L and P270L.

### Screening with 15 amino acid overlapping peptides to identify MHC-II epitopes in Cda2 and Cda1

A more rigorous screening approach was taken to validate the hits identified with the long peptides and to identify epitopes that may have been missed. Long peptides require uptake, processing, and presentation by APCs whereas 15mers can directly bind in the MHC-II pocket where they can be presented to the TCR on CD4^+^ T cells (14–16). We had peptide libraries synthesized spanning Cda2 (72 peptides) and Cda1 (71 peptides) that consisted of 15mers overlapping by 10 amino acids. DR4 and C57BL/6 mice were immunized with either Cda2 or Cda1 adjuvanted with CAF01, spleens were harvested, and splenocytes were stimulated with the peptides from the respective Cda libraries. IFNγ levels in the supernatants served as the readout for immune stimulation. When a hit was obtained, the epitope was precisely mapped using 15mers that overlapped by 14 amino acids. For each cryptococcal protein and MHC-II allele, the IEDB was used to generate the predicted IC50 values for each 15mer within the protein (22) and plotted against the position of the peptide in the protein sequence. The IC50 is the nM concentration of peptide required to displace a reference peptide from the MHC-II molecule (https://iedb.org/home_v3.php). The lower the IC50, the stronger the binding. While minimum binding thresholds have not been well established and differ for each MHC-II allele, nearly all validated MHC-II epitopes reportedly have IC50 values under 1,000 nM (https://help.iedb.org/hc/en-us/articles/114094151811-Selecting-thresholds-cut-offs-for-MHC-class-I-and-II-binding-predictions).

The DRB1*0401 epitopes identified using DR4 mice immunized with Cda2 were mapped first (Figure 2). Figure 2A shows the IFNγ levels stimulated by each of the 15mers from the Cda2 peptide library. The predicted IC50 values are superimposed on the graph. Hits were obtained for peptides P35, P50, P55, P70, and P75. Each of the 15mer sequences is also found in a 30mer: P35 in P20L; P50 and P55 in P45L; and P70 and P75 in P60L and P70L. Both screens validated that the DRB1*0401 MHC-II epitopes are confined to the N-terminal region of Cda2. We next mapped each of the epitopes using 15mers that overlapped by 14 amino acids. With this mapping, we found an epitope that included P35 (Figure 2B), one that included P50 and P55 (Figure 2C), and a third one that included P70 and P75 (Figure 2D). Thus, Cda2 contains three distinct epitopes that stimulate splenocytes from Cda2-immunized DR4 mice. A 9-amino acid core sequence for each epitope was predicted by NetMHCIIpan 4.1 BA software as being shared among the epitope-defining peptides that were stimulatory. The three distinct epitopes within Cda2 were named P36, P58, and P68 based on the P# of the peptide containing the 9-amino acid sequence that elicited high IFNy levels while having a low predicted IC50.

**Figure 2.**
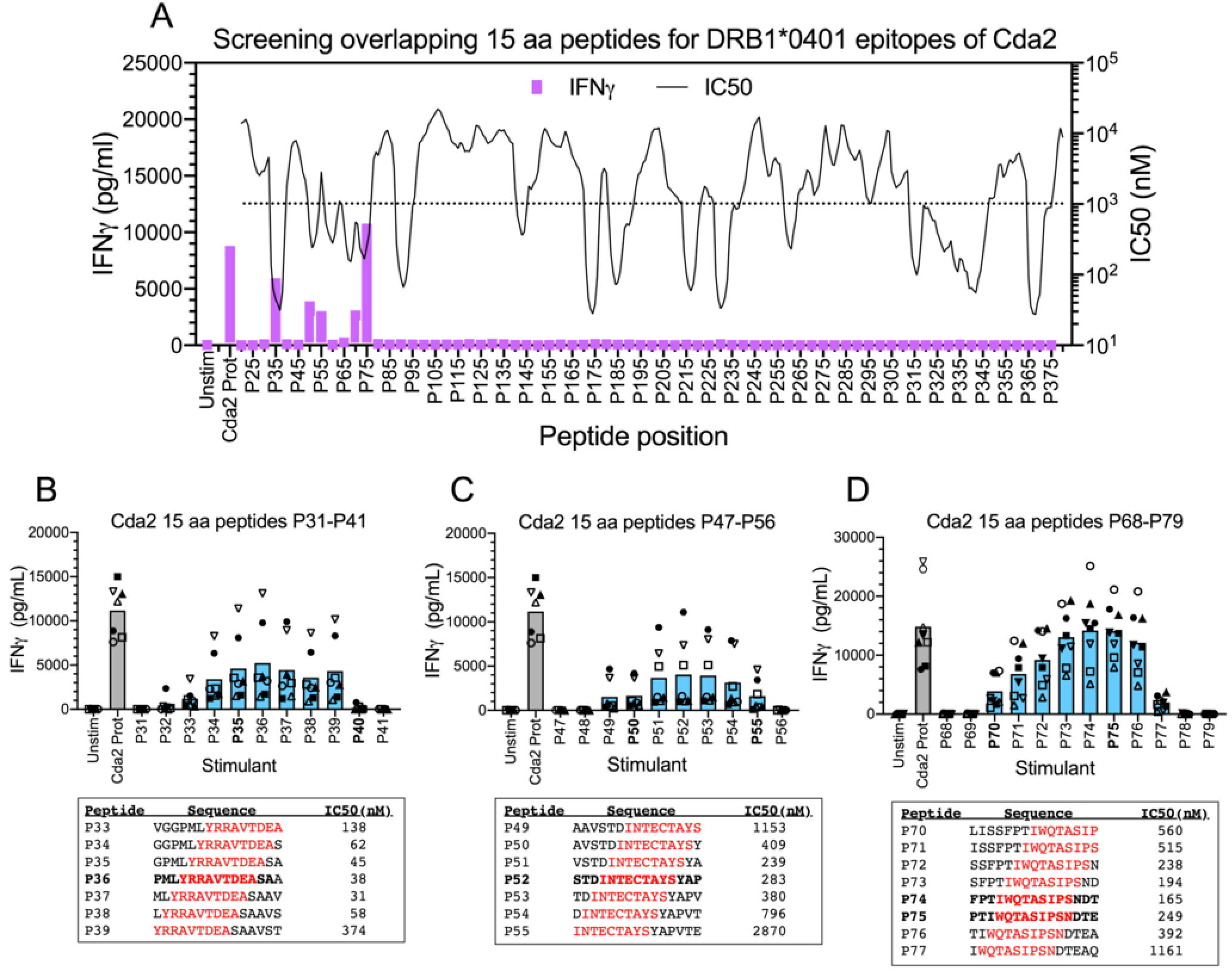
Mapping DRB1*0401 epitopes of Cda2 using 15 aa overlapping peptides. As in Figure 1 except splenocytes from DR4 mice vaccinated with recombinant Cda2 protein (Cda2 Prot) were assayed using 15 aa peptides as stimulants. (**A**) Screening the Cda2 recombinant protein with overlapping 15 aa peptides (n=3 mice). Average IFNγ generated by each stimulant is represented by the height of its respective purple bar. Predicted IC50 values (nM) for sequential 15 aa peptides of recombinant Cda2 protein were plotted as a solid line. The dotted horizontal line is set at 1,000 nM as most validated MHC-II epitopes have IC50 values under this value. (**B-D**) Three epitopes discovered in **(A)** were mapped using splenocytes from immunized mice stimulated with 15 aa peptides which overlap by 14 aa. Below each graph are listed the sequences of peptides that stimulated IFNγ generation and their respective IC50 value predicted for binding to the human DRB1*0401 MHC-II allele. The shared 9 aa core sequence of each epitope is in red type. The peptide with the 9 aa core positioned in the middle of the peptide is in bold. Data are from two independent experiments, each with 3-4 mice. Each symbol shape represents results from an individual mouse. Closed symbols are from the first experiment whereas open symbols are from the repeat experiment. Statistical comparisons are shown in Table S3A.

We next mapped the IAb MHC-II epitopes in C57BL/6 mice immunized with Cda2 (Figure 3). Although the predicted IC50 values revealed relatively few potential epitopes with threshold values less than 1,000 nM (Figure 3A), the long peptides P20L, P45L, P60L and P70L, and 15mers P35 and P70 identified epitopes in the same region as those mapped using DR4 mice. Fortunately, we were able to repurpose the 15mers that overlap by 14 amino acids to do the fine mapping. Three IAb MHC-II epitopes were also discovered in this region. The epitope mapped in Figure 3B is shared with its DRB1*0401 counterpart and has the same 9 amino acid core. The epitope in Figure 3C is found within the long peptide P45L but was not identified in the screen with the 15mers. The epitope in Figure 3D shares a stimulatory peptide, P70 with the DRB1*0401 epitope shown in Figure 2C. Three additional epitopes were identified in the screens using C57BL/6 mice and were mapped in Figure 3E-G. The epitope in Figure 3E was identified with long peptides P203L, P220L and with 15mers P215 and P220. This epitope appears to generate the dominant IFNy response. The epitope in Figure 3F is near the one mapped in Figure 3E; its 15mers are also embedded in the 30mer peptide P220L but was clearly distinguished in the 15mer screen by stimulation of P230. The epitope in Figure 3G was identified in the screens by 30mer P295 and 15mer P310. Interestingly, of the six epitopes that were discovered and their 9-amino acid core sequences identified (Figure 3B-G), three of these IAb epitopes had predicted IC50 values above 1,000 nM.

**Figure 3.**
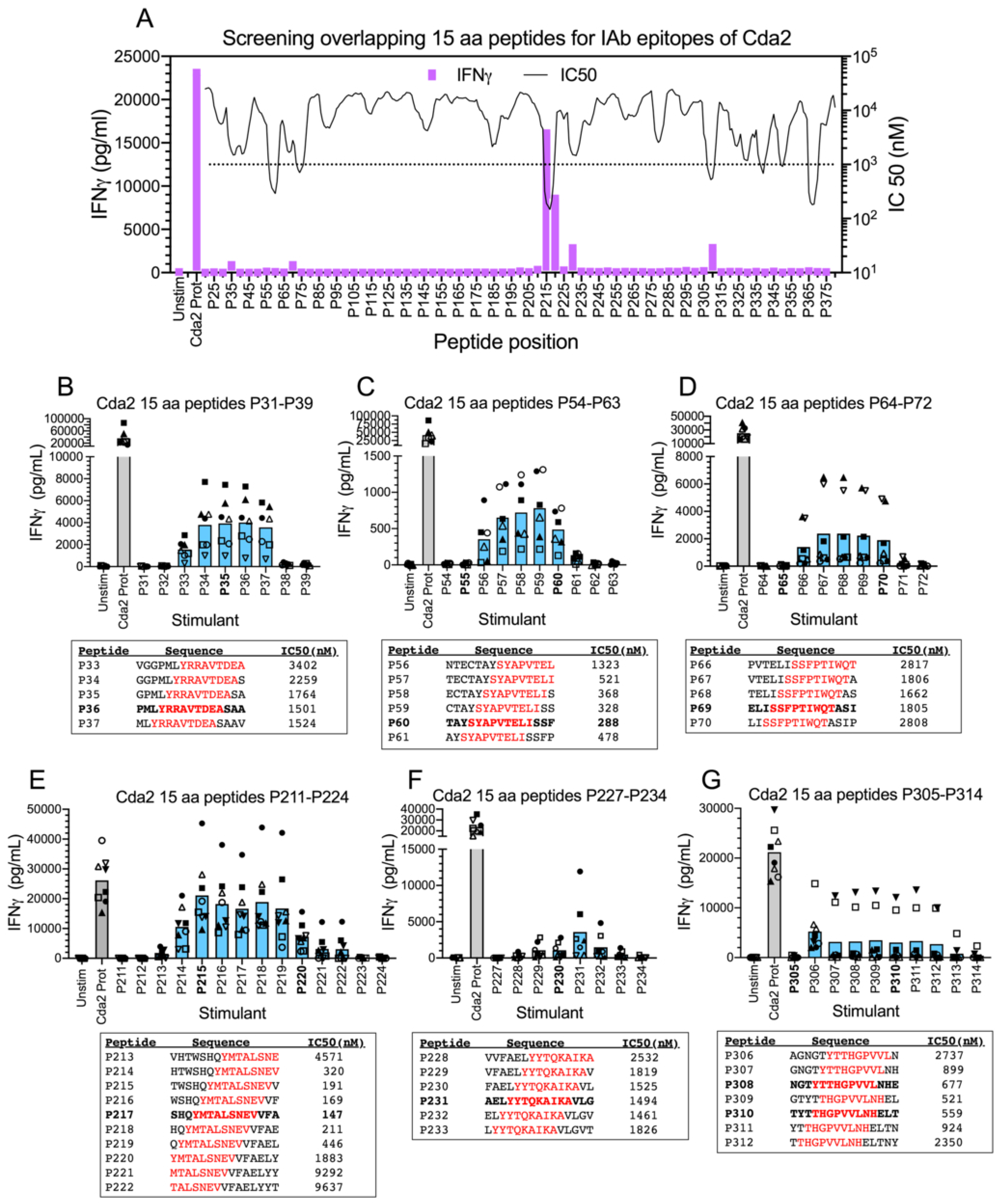
Mapping IAb epitopes of Cda2 using 15 aa overlapping peptides. As in Figure 2 except splenocytes from C57BL/6 mice vaccinated with recombinant Cda2 protein (Cda2 Prot) were assayed. (**A**) Screening the Cda2 recombinant protein with overlapping 15 aa peptides (n=3 mice). Averaged IFNγ generated by each stimulant is represented by the height of its respective purple bar. Predicted IC50 values (nM) for sequential 15 aa peptides of recombinant Cda2 protein were plotted as a solid line. The dotted horizontal line is set at 1,000 nM as most validated MHC-II epitopes have IC50 values under this value. (**B-G**) Six epitopes were mapped using splenocytes from immunized mice stimulated with 15 aa peptides which overlap by 14 aa. Below each graph are listed the sequences of peptides that stimulated IFNγ generation and their respective IC50 value predicted for binding to the mouse IAb MHC-II allele. The shared 9 aa core sequence of each epitope is in red type. The peptide with the 9 aa core positioned in the middle of the peptide is in bold. Data are from two independent experiments, each with 3-4 mice. Each symbol shape represents results from an individual mouse. Closed symbols are from the first experiment whereas open symbols are from the repeat experiment. Statistical comparisons are shown in Table S3B.

MHC-II epitopes in DR4 and C57BL/6 mice immunized with Cda1 were determined using the same methodology used for discovering Cda2 epitopes. Consistent with the screening performed with the long peptides, two distinct DRB1*0401 epitopes were validated in immunized DR4 mice using the overlapping 15mers screen (Figure 4A). The sequence of the 15mer P325 is in the sequence of the 30mer P320L and the sequence of 15mer P355 within the 30mer P345L. The mapping of each epitope and identification of their respective 9 amino acid core sequences are shown in Figure 4B and 4C, respectively. Two IAb epitopes were also validated using vaccinated C57BL/6 mice (Figure 5). The sequence of the 15mer P135 is in 30mer P120L and that of 15mer P270 is in 30mer P270L. Mapping of each epitope and identification of their respective 9 amino acid core sequences are shown in Figure 5B and 5C, respectively. The sequences of the stimulating peptides of the epitope mapped in Figure 5B contain three cysteines, which is unusual. The predicted IC50 values for the DRB1*0401 epitopes were low (Figure 4B and 4C). However, for the C57BL/6 mice, the IAb IC50 values were very high (Figure 5B and 5C), even exceeding >10,000 nM for the epitope mapped in Figure 5B.

**Figure 4.**
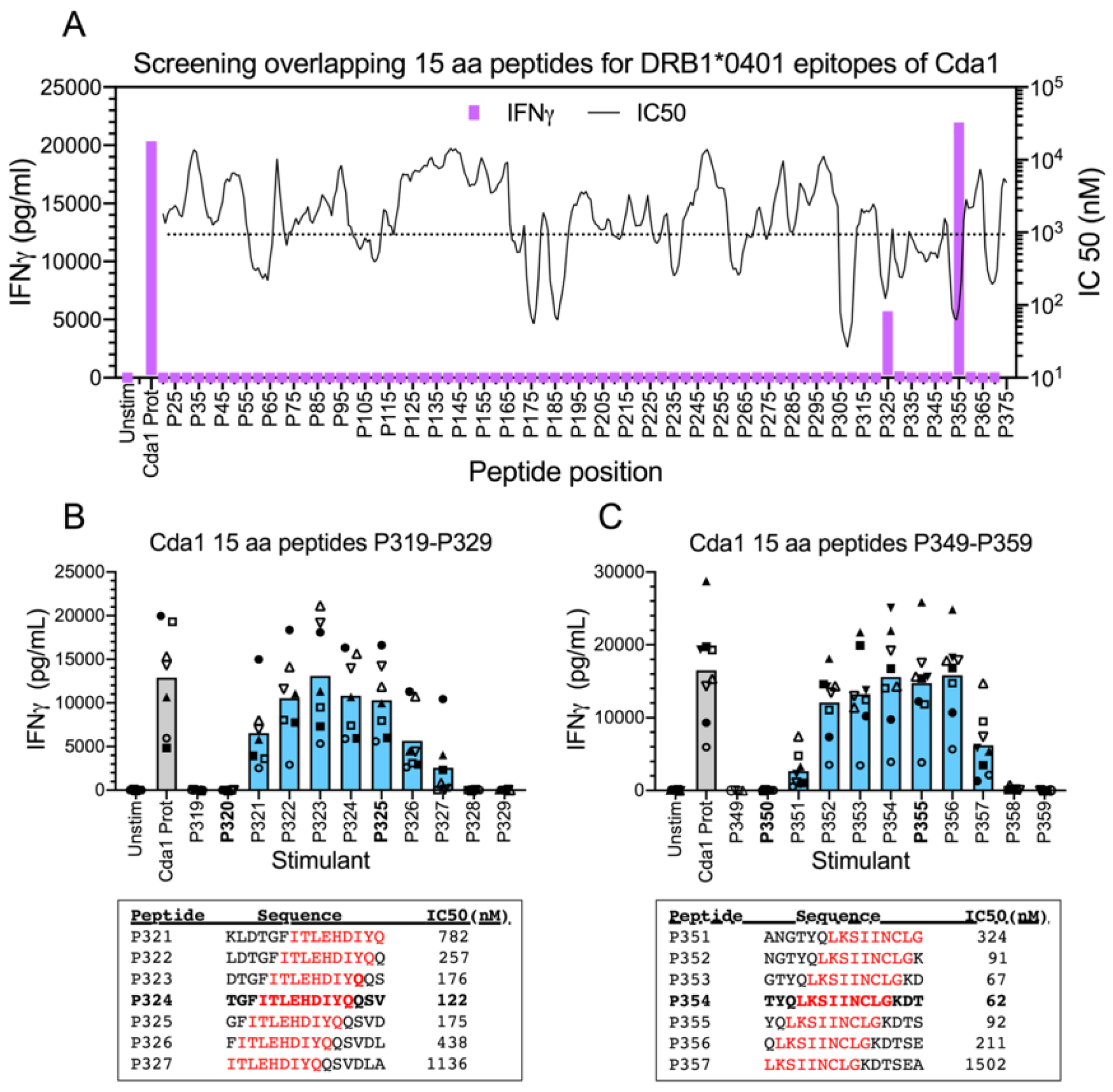
Mapping DRB1*0401 epitopes of Cda1 using 15 aa overlapping peptides. As in Figure 2 except splenocytes from DR4 mice vaccinated with recombinant Cda1 protein (Cda1 Prot) were assayed. (**A**) Screening the Cda1 recombinant protein with overlapping 15 aa peptides (n=3 mice). Averaged IFNγ generated by each stimulant is represented by the height of its respective purple bar. Predicted IC50 values (nM) for sequential 15 aa peptides of recombinant Cda1 protein were plotted as a solid line. The dotted horizontal line is set at 1,000 nM as most validated MHC-II epitopes have IC50 values under this value. (**B,C**) Two epitopes were mapped using splenocytes from immunized mice stimulated with 15 aa peptides which overlap by 14 aa. Below each graph are listed the sequences of peptides that stimulated IFNγ generation and their respective IC50 value predicted for binding to the human DRB1*0401 MHC-II allele. The shared 9 aa core sequence of each epitope is in red type. The peptide with the 9 aa core positioned in the middle of the peptide is in bold. Data are from two independent experiments, each with 3-4 mice. Each symbol shape represents results from an individual mouse. Closed symbols are from the first experiment whereas open symbols are from the repeat experiment. Statistical comparisons are shown in Table S4A.

**Figure 5.**
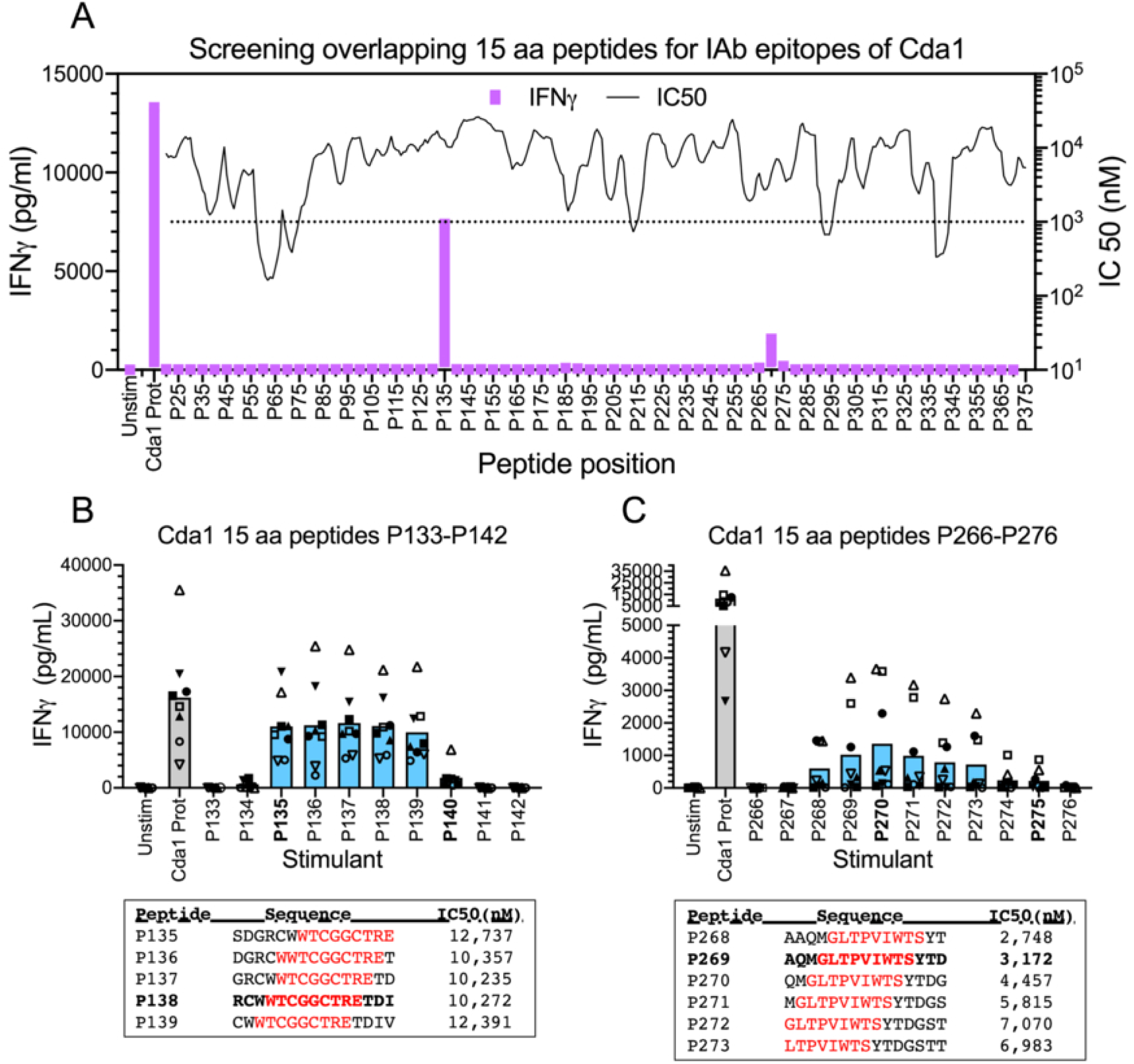
Mapping IAb epitopes of Cda1 using 15 aa overlapping peptides. As in Figure 2 except splenocytes from C57BL/6 mice vaccinated with recombinant Cda1 protein (Cda1 Prot) were assayed using 15 aa peptides as stimulants. (**A**) Screening the Cda1 recombinant protein with overlapping 15 aa peptides (n=3 mice). Averaged IFNγ generated by each stimulant is represented by the height of its respective purple bar. Predicted IC50 values (nM) for sequential 15 aa peptides of recombinant Cda1 protein were plotted as a solid line. The dotted horizontal line is set at 1,000 nM as most validated MHC-II epitopes have IC50 values under this amount. (**B,C**) Two epitopes were mapped using splenocytes from immunized mice stimulated with 15 aa peptides which overlap by 14 aa. Below each graph are listed the sequences of peptides that stimulated IFNγ generation and their respective IC50 value predicted for binding to the mouse IAb MHC-II allele. The shared 9 aa core sequence of each epitope is in red type. The peptide with the 9 aa core positioned in the middle of the peptide is in bold. Data are from two independent experiments, each with 4 mice. Each symbol shape represents results from an individual mouse. Closed symbols are from the first experiment whereas open symbols are from the repeat experiment. Statistical comparisons are shown in Table S4B.

In summary, the identification and mapping of the nine epitopes of Cda2 and four epitopes of Cda1 has shown that these MHC-II epitopes span 5-9 sequential 15 amino acid peptides that generate IFNy above the background levels of flanking peptides. For each of the epitopes we identified, statistical analysis by ordinary one-way ANOVA was done using pairwise comparisons of Unstim IFNy values to those that were peptide-stimulated (Tables S3 and S4). P values of peptides of stimulatory peptides were lower than the non-stimulatory flanking peptides.

Effect of CD4^+^ T cell depletion on IFNγ release stimulated by validated epitopes. The adjuvant we used, CAF01, stimulates strong Th1- and Th17-biased CD4^+^ T cell responses (8, 26). However, we have also observed weak Th1-biased CD8^+^ T cell responses following CAF01-adjuvanted vaccination (8). Therefore, we next determined if CD4^+^ T cells were the source of the IFNγ following stimulation of splenocytes from immunized DR4 and C57BL/6 mice for validated epitopes of Cda2 and Cda1. To accomplish this, we compared IFNγ release in splenocytes depleted of CD4^+^ T cells with mock-depleted splenocytes. Efficiency of depletion was verified by flow cytometry (Figure S2 and Table S5). Unstimulated cells served as the negative control while cells stimulated with 12-phorbol 13-myristate acetate (PMA) plus ionomycin (IO) served as the positive control. Regardless of the mouse strain and the protein used for immunization, peptide-stimulated IFNγ levels of each epitope were reduced to background concentrations when CD4^+^ T cells were depleted (Figure 6A-D). The cells retained the capacity to make IFNγ, as the PMA+IO stimulated levels of IFNγ were roughly the same when comparing CD4-depleted to mock-depleted splenocytes. Thus, CD4^+^ T cells are the apparent source of the epitope-stimulated IFNγ release from splenocytes of Cda2 and Cda1 immunized mice.

**Figure 6.**
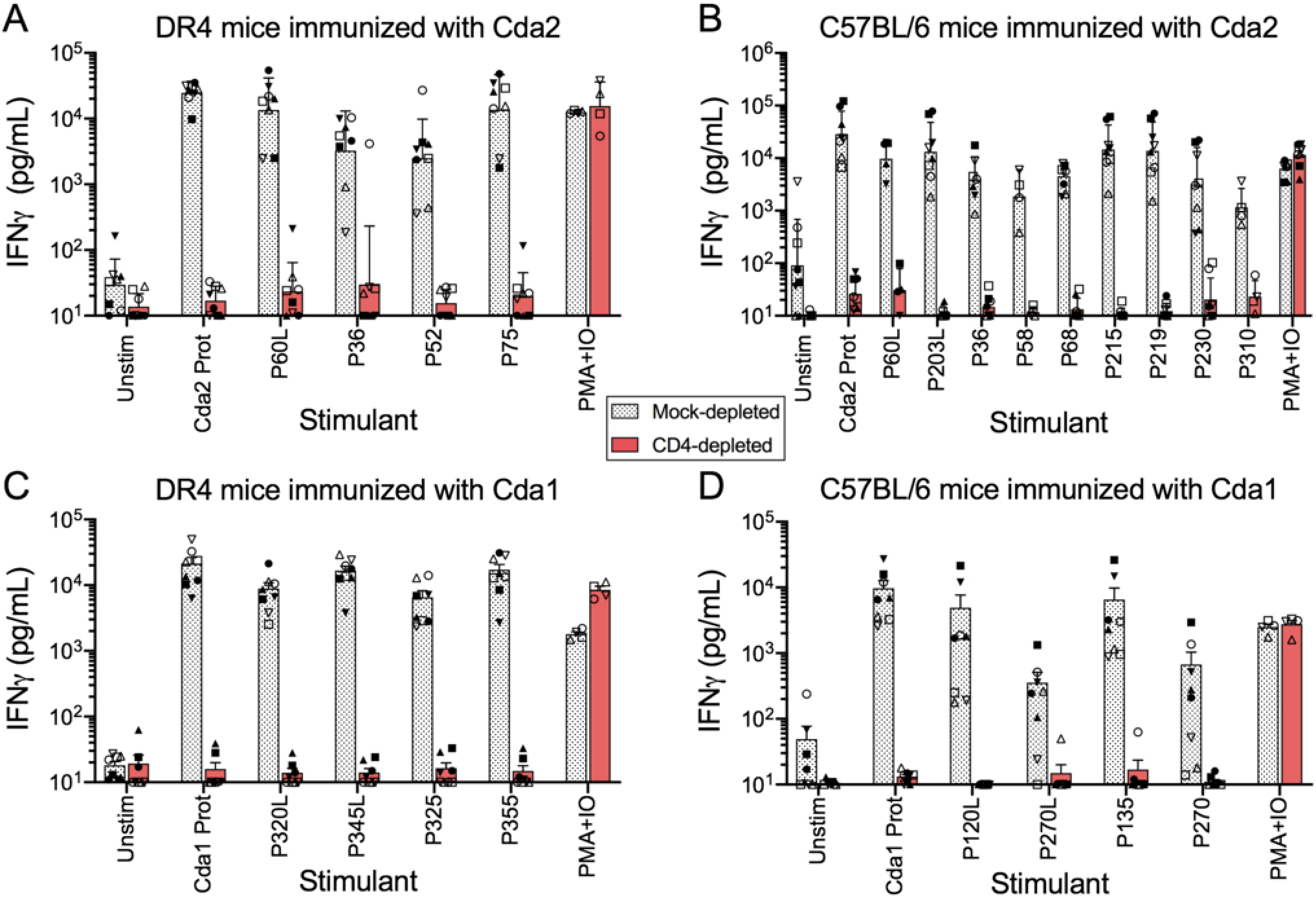
The effect of CD4^+^ T cell depletion on splenocyte stimulation by Cda1 and Cda2 epitopes. DR4 and C57BL/6 mice were immunized with either Cda2 (**A,B**) or Cda1 (**C,D**) proteins, respectively. Spleens were harvested two weeks after the last vaccination and single cell suspensions were prepared. The splenocytes were then either mock-depleted or depleted of CD4^+^ T cells using magnetic beads (CD4-depleted). The mock-depleted group was submitted to the same conditions except no beads were added. The two splenocyte populations were either left unstimulated (Unstim) or stimulated with the indicated protein (Prot) or peptide. Phorbol 12-myristate 13-acetate + ionomycin (PMA+IO) was used as a positive control. Cells were cultured for three days following which IFNγ levels in the supernatant were measured by ELISA. Data are from two independent experiments, each with 4 mice. Each symbol shape represents results from an individual mouse. Closed symbols are from the first experiment whereas open symbols are from the repeat experiment.

### Construction of a Cda2/1 hybrid protein and its testing in a vaccine model

For DR4 mice, the three validated epitopes (P36, P52, and P75, shown in Figure 2) in Cda2 are near the N-terminus while both validated epitopes of Cda1 (P324 and P354, shown in Figure 4) are near the C-terminus (Figures 2 and 4). We next constructed a hybrid protein in *E. coli*, which we named Cda2/1, by combining amino acids 29-95 of Cda2 with amino acids 316-374 of Cda1 (Figure S3). The DRB1*0401 and IAb binding profiles, based on predictions using the IEDB, are shown in Figure 7A and 7B, respectively. Next, splenocyte preparations were made from DR4 mice immunized with Cda2/1 and then incubated with P36, P52, P75, P324, and P354. Each of the five peptides stimulated IFNγ release, as did full length Cda2, Cda1, and Cda2/1 hybrid protein (Figure 7C). IFNγ secretion was not seen following stimulation with two peptides (designated Hyb-P58 and Hyb-P63), which contained amino acid sequences from both Cda2 and Cda1.

**Figure 7.**
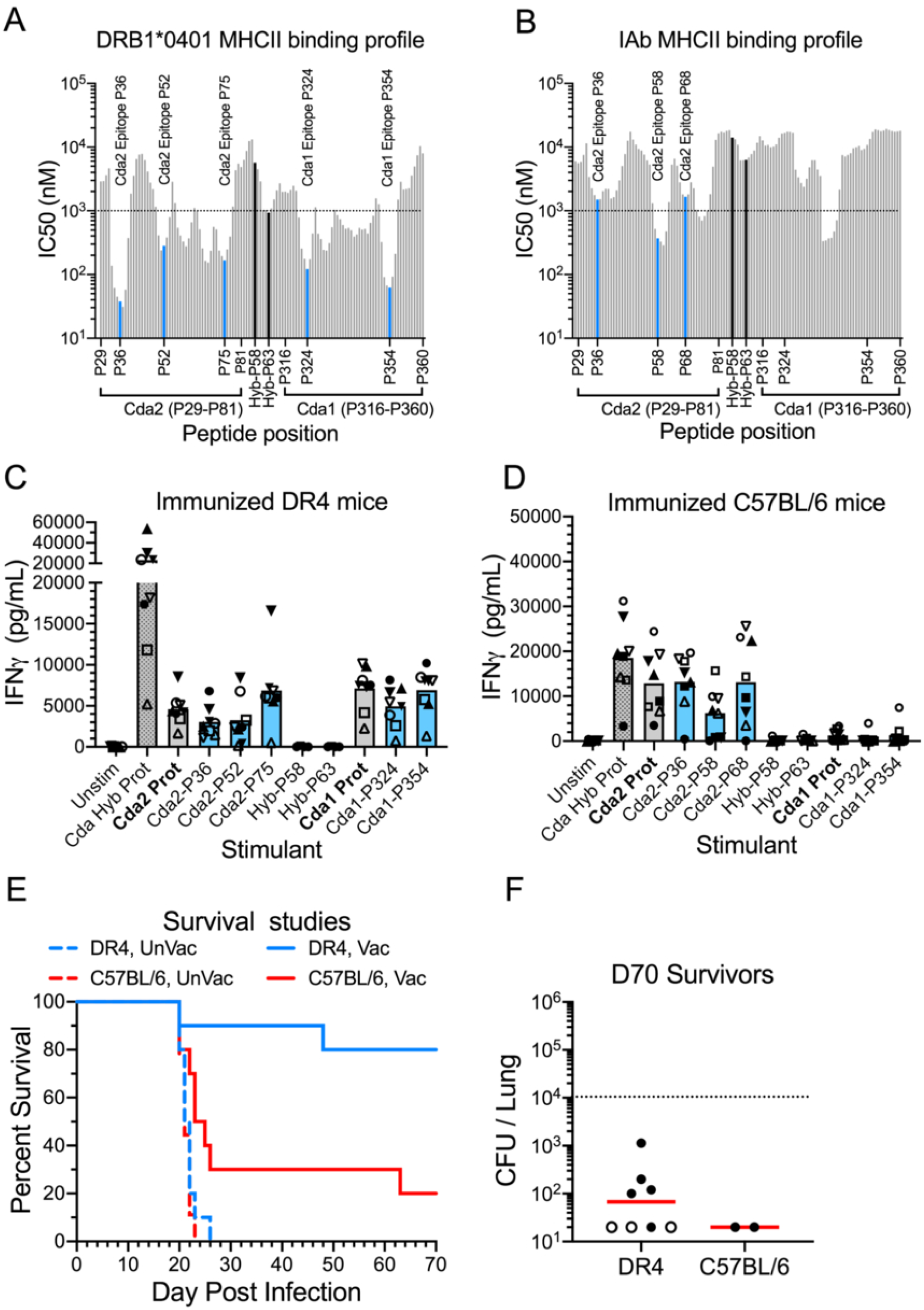
Analysis of the Cda2/1 hybrid protein as an immunogen and vaccine antigen. (**A,B**). IC50 profiles of the sequential 15 aa peptides that comprise the Cda2/1 protein for MHC-II alleles HLA-DRB1*0401 (DR4 mice, in [**A**]) and H2-IAb (C57BL/6 mice in [**B**]), respectively. Blue bars denote the location of 15 aa peptide sequences which stimulated IFNγ in immune splenocytes. The black bars show the location of 15 aa peptides (Hyb-P58 and Hyb-P63) containing aa sequences present in Cda2 and Cda1 (see Figure S3). The dotted horizontal line is set at 1,000 nM as most validated MHC-II epitopes have IC50 values under this amount. (**C,D**). Splenocytes from DR4 (**C**) and C57BL/6 (**D**) mice immunized with the Cda2/1 hybrid protein were stimulated with the 15 aa peptides identified with the blue and black bars in (**A**) and (**B**). The negative and positive controls were unstimulated splenocytes (Unstim) and splenocytes stimulated with recombinant Cda2/1 protein (Cda Hyb Prot). After three days incubation, IFNγ was measured by ELISA in supernatants of splenocyte cultures. Data are from two independent experiments, each with 4 mice. Each symbol shape represents results from an individual mouse. Closed symbols are from the first experiment whereas open symbols are from the repeat experiment. (**E,F**). DR4 and C57BL/6 mice were left unvaccinated (UnVac) or vaccinated (Vac) with Cda2/1 and then given a pulmonary challenge with *C. neoformans*. Survival curves are shown in (**E**) and represent data from two independent experiments, each with 5 mice per group. P<0.01 comparing survival curves of vaccinated DR4 and C57BL/6 mice. P<0.0001 comparing UnVac and Vac mice for both DR4 and C57BL/6 mice. Lung CFUs of mice that survived to D70 are shown in (**F**). Closed and open circles represent CFUs of surviving mice from the first and second experiments, respectively. The dotted horizontal line at 10^4^ CFUs denotes the challenge *C. neoformans* inoculum.

For the C57BL/6 mice, only the P36, P58, and P68 IAb epitopes of Cda2 (mapped in Figure 3A-C) were retained in Cda2/1. The three other validated IAb epitopes in Cda2 and the two validated IAb epitopes of Cda1 were not included in the Cda2/1 construct. As expected, P36, P58, and P68 stimulated IFNγ production in splenocytes obtained from C57BL/6 mice immunized with Cda2/1 (Figure 7D). Peptides Hyb-P58 and Hyb-P63 and peptides containing sequences from the Cda1 portions of Cda2/1did not stimulate. When the recombinant proteins were used as stimuli, the Cda2 and hybrid Cda2/1 proteins, but not the Cda1 protein, induced IFNγ production. This result was consistent with the absence of Cda1 IAb epitopes in the Cda2/1 construct.

In the final set of experiments, we vaccinated DR4 and C57BL/6 mice with Cda2/1 adjuvanted with CAF01. Mice received a prime vaccine followed three weeks later by a booster dose. Both mouse strains were protected from *C. neoformans* challenge by the Cda2/1 vaccine, with protection significantly greater in the DR4 mice (Figure 7E). Many of the surviving mice had undetectable lung CFUs (Figure 7F).

## Discussion

Multiple lines of evidence support the inclusion of *C. neoformans* Cda2 and Cda1 in subunit cryptococcal vaccine formulations. Cda2 and Cda1 protein sequences lack human homologs, are abundantly expressed during cryptococcal infection, are highly conserved among cryptococcal species, and elicit protective and immunodominant Th1-biased CD4^+^ T cell responses in murine models of cryptococcosis (6–11, 27, 28). However, to be effective, CD4^+^ T cell vaccines need to account for the extensive variation in MHC-II molecules expressed in the human population (16). Here, we used experimental data, supported in part by in silico predictions to map the MHC-II epitopes in Cda2 and Cda1 that stimulate CD4^+^ T cell responses in C57BL/6 and DR4 mice. We then designed a hybrid protein containing validated epitopes of Cda2 and Cda1 to demonstrate that an adjuvanted vaccine containing the hybrid protein could protect humanized mice against an otherwise lethal cryptococcal infection.

The lower the predicted IC50, the tighter the predicted binding of a peptide to an MCH-II molecule, with a threshold of an IC50 < 1,000 nM recommended to predict a good binder (29, 30). Using this cutoff, most 15mer peptides present in Cda2 and Cda1 that were predicted by the IEDB to be good binders to DRB1*04:01 nevertheless did not stimulate IFNγ responses in splenocytes from immunized mice. An antigen-stimulated CD4^+^ T cell response requires antigen-processing and presentation of the peptide to the TCR. Even assuming the accuracy of the bioinformatic predictions, loss of epitopes can occur if antigen-processing does not lead to MHC-II presentation of the peptide. This could occur during processing by cathepsins or if posttranslational modifications, such as glycosylation, interferes with presentation (23). Moreover, even if peptide binding to MHC-II is strong, the precursor frequency of TCRs capable of binding to the T cell exposed motif on the peptide could be insufficient to initiate a CD4^+^ T cell response (31). Regardless of the reasons why a predicted MHC-II binding epitope is not validated, these data emphasize the importance of verifying in silico predictions experimentally.

Unexpectedly, we also validated epitopes that were not predicted to bind well to MHC-II. For example, the Cda1 P135 peptide stimulated robust IFNγ production in splenocytes from immunized C57BL/6 mice despite having a predicted IC50 of >10,000 nM to H2-IAb. This epitope would not have been discovered had we not taken an unbiased approach to epitope discovery by screening libraries of overlapping peptides rather than just examining peptides with predicted IC50 values of <1,000 nM. The possibility that the IFNγ was from cells other than CD4^+^ T cells was effectively eliminated, as IFNγ production was reduced to background levels after ex vivo depletion of CD4^+^ T cells from splenocytes. It is important to note, however, that our experimental setup, including the use of CAF01 as an adjuvant and screening with 15mer peptides, was designed to detect MHC-II epitopes. Immunization with an adjuvant that stimulates strong CD8^+^ T cell responses and then screening with 9mer peptides could validate which sequences are MHC-I epitopes.

Not surprisingly considering the evolutionary distance between mice and humans, most peptides that were validated epitopes in DR4 mice were not stimulatory in C57BL/6 mice. About 10% of humans express at least one DRB1*0401 allele (20). The highly polymorphic nature of human MHC-II (13, 19, 20) raises the question of whether epitopes that bind to DRB1*0401 and stimulate subsequent CD4^+^ T cell responses will do the same with other DRB1 molecules. While this needs to be determined experimentally, IEDB predictions suggest that the epitopes we validated in the DR4 mice will bind to other (although not all) DRB1 molecules (Table S6). Ultimately, to be protective, we postulate that CD4^+^ T cell vaccines will need to account for the extensive diversity of human MHC-II alleles by containing multiple stimulatory epitopes. Experiments examining transgenic mice expressing humanized DRB1 molecules other than DRB1*0401 are planned. Contributions of other MHC-II molecules also need to be considered given studies suggesting HLA MHC-II DQB and DPB genotypes are also associated with risk for human cryptococcosis (32, 33). Finally, our data emphasize the importance of taking into account MHC-II differences between mice and humans when developing T cell vaccines using murine models.

We initially tested peptide libraries consisting of overlapping long peptides, ≥30 amino acids in length, but then switched to overlapping 15mers. Compared with 15mers, screening 30mers requires fewer peptides given the longer length of the peptide. However, for positive hits, the mapping of the epitope to identify the core 9 amino acid sequence that binds to MHC-II and is exposed to T cells (17, 18) may require approximately twice the number of peptides when the primary screen is with long peptides. Additionally, as noted above, 15mers can directly bind in the MHC-II pocket, bypassing the need for antigen uptake and processing by APCs (14–16). Perhaps most importantly, when screening overlapping peptides, sequential peptides should have sufficient overlap so that epitopes are not missed. For the 15mers, we chose an overlap of 10 amino acids which should be sufficient for detecting nearly all epitopes given that we experimentally showed that an epitope is defined by 5-9 stimulatory 15mers and the 9 amino acid core region is shared by the stimulatory peptides (17, 18).

Variations in reactivity (as measured by peptide-stimulated splenocyte IFNγ production) among individual mice used as experimental replicates were observed. Generation of T cell receptors occurs via stochastic processes which can result in large differences in individual TCR repertoires even in inbred mouse strains (29). The relative contribution of TCR precursor frequencies to these variations seen in our mouse experiments is speculative though. The translational implication is that cryptococcal T cell vaccines need to contain multiple epitopes to not only account for population diversity in HLA alleles but also for individual variations in TCR precursor frequencies.

Epitope mapping has translational relevance as it informs construction of multi-epitope hybrid proteins that can be used in subunit T cell vaccines and possibly diagnostic testing. From a manufacturing standpoint, a single protein vaccine is less expensive to manufacture and test than one containing multiple proteins. In addition, eliminating portions of the proteins that are not contributing to protection decreases the potential for eliciting autoimmunity. As a proof-of-principle, we constructed a relatively small (133 amino acid) hybrid protein incorporating the epitopes in Cda2 and Cda1 that stimulated CD4^+^ T cell responses in DR4 mice immunized with these two proteins. DR4 mice vaccinated with the hybrid protein adjuvanted with CAF01 were robustly protected against an otherwise lethal cryptococcal pulmonary challenge. In contrast, C57BL/6 mice were only minimally protected, which was not surprising as the hybrid vaccine protein included only three of the six identified Cda2 epitopes and none of the Cda1 CD4^+^ T cell epitopes for the MHC-II of that mouse strain. Nevertheless, the hybrid protein still elicited strong ex vivo splenocyte IFNγ responses in immunized C57BL/6 mice; this response likely accounted for the protection that was observed.

A caveat to using hybrid proteins in vaccines is they can create neoantigens (also known as junctional epitopes) at fusion sites. In future studies, this potential pitfall may be mitigated by adding linkers or employing bioinformatics to eliminate neoantigens with undesirable properties such as possible human homology (34, 35). In addition to Cda2 and Cda1, we have discovered other proteins that, when formulated as adjuvanted vaccines, protect mice against otherwise lethal cryptococcal infections (10, 11). Epitope mapping of these proteins is underway. Once completed, we plan to create hybrid vaccines that contain multiple epitopes that elicit strong IFNγ responses. The protective efficacy of such vaccines will be tested using clinical strains representative of the global lineages of *C. neoformans* and *C. gattii* (7, 36), and in strains of mice transgenic for human MHC-II. Ultimately though, vaccines will need to advance from preclinical studies to human testing, including in the immunocompromised populations most in need of a cryptococcal vaccine.

## Materials and Methods

### Mice

Two strains of mice were used: C57BL/6 (Strain 000664; The Jackson Laboratory, Bar Harbor, ME) and DR4 (Strain 4149; Taconic Biosciences, Rensselaer, NY). Mice were maintained at University of Massachusetts Chan Medical School (UMCMS) and housed in a pathogen-free facility. Experimental procedures were approved by the UMCMS Institutional Use and Care of Animals Committee.

### Immunizations and vaccinations

*C. neoformans* proteins Cda2 (CNAG_01230) and Cda1 (CNAG_05799) were cloned into plasmid pET19b to position a 10x His tag at the N-terminus of each protein (11). The DNA sequence encoded amino acids 20-378 of Cda2 and amino acids 20-374 of Cda1. The first 19 amino acids were not included as they contain the cleaved signal sequences of the two proteins. The Cda2/1 hybrid protein was generated by coupling DNA encoding amino acids 29-95 of Cda2 to amino acids 316-374 of Cda1. The DNA sequence encoding the hybrid protein was synthesized by Genscript (Piscataway, NJ) and cloned into vector pET21b to include a 6x His tag at the C-terminus. Plasmids were transformed into *E. coli* strain BL21 DE3.

Recombinant proteins were expressed in *E. coli* strain BL21 DE3 using Overnight Express Instant TB medium, 1% glycerol (EMD Millipore, Burlington, MA); cultures were shaken at 30°C for 18-24 h. Extraction of protein from *E. coli* cell pellets and purification of recombinant protein on His-Bind Resin (EMD Millipore) were done in buffers containing 6 M urea, as previously described (11). Stocks of recombinant protein at 5 mg/mL in 6 M urea, 20 mM Tris, pH 7.9 were stored at −80°C. Each dose of vaccine (100 µL) was formulated to contain 5 µg of protein diluted in Tris buffer with the liposomal-based CAF01 (6–8) as the adjuvant. CAF01 (Statens Serum Institut, Copenhagen, Denmark) contains 2,500 μg/mL N,N’-dimethyl-N,N’-dioctadecylammonium and 500 μg/mL α,α′-trehalose 6,6′-dibehenate in Tris Buffer (10 mM, pH 7.0) supplemented with 2% glycerol. For immunization of the mice, two 100 µL subcutaneous injections were done, one injection into each side of the abdomen, and spaced about 1 cm apart. The second immunization was given three weeks later. Mice were euthanized two weeks after the second immunization and spleens were harvested.

Vaccine efficacy was determined as in our previous studies (7, 8). Briefly, groups of five mice were left unvaccinated or vaccinated as above. Two weeks after the second vaccination, mice were anesthetized with 2% isoflurane (Pivetalvet) and challenged with *C. neoformans* strain KN99α (37) by orotracheal infusion of 1 x 10^4^ CFU into the lungs. Following infection, the mice were monitored daily. Mice that survived 70 days post infection (DPI) were euthanized. Lungs were homogenized in 4 mL phosphate buffered saline (PBS) with 200 U/mL penicillin and 200 µg/mL streptomycin and plated on Sabouraud dextrose agar. CFUs were counted following incubation at 30°C. Detection limits were 20 CFU/lung.

### Peptides

Peptides were synthesized by Genscript. Long peptides 30-35 amino acids in length exceeded 80% purity and 15 amino acid peptides exceeded 70% purity. Peptides were provided as lyophilized powder in vials containing either 2.0 mg or 0.5 mg of peptide. Each peptide was dissolved with 100% dimethyl sulfoxide (#BP231-100, Fisher Scientific) to generate stock solutions at 5 mg/mL, which were stored at −20°C.

### Splenocyte stimulation and IFNγ ELISA

Splenocytes were stimulated with proteins and peptides as described (6, 7). Briefly, spleens were homogenized in complete media (RPMI 1640 supplemented with 10% FBS, 1% HEPES, 1% GlutaMAX, and 1% Penicillin–Streptomycin; reagents purchased from Thermo Fisher Scientific) using the piston of a 3 mL syringe on a 70 µm cell strainer. The cell pellet was collected following centrifugation, red blood cells were lysed, and the splenocytes were washed twice in complete media. The concentration and viability of cells were determined with Trypan blue (Bio-Rad Laboratories, Hercules, CA) using a T20 cell counter (Bio-Rad). Splenocytes were plated in tissue culture-treated 96-well round bottom plates (#3799, Corning Inc, Corning, NY) at 1 × 10^6^ cells/well. Cells were left unstimulated or stimulated with recombinant protein (5 µg/mL) or peptide (2.5 µg/mL). Duplicate wells were made for each stimulant. Incubations were at 37°C in humidified air supplemented with 5% CO2. Splenocytes were cultured for three days in complete media. Plates were then centrifuged, supernatants were collected and stored at −80°C. IFNγ concentrations in thawed supernatants were determined with the R&D Systems Mouse IFNγ DuoSet ELISA Kit (Bio-Techne, Minneapolis, MN) according to the manufacturer’s instructions. Samples were diluted 1:2, or if needed 1:10. The lowest standard was 10 pg/mL. Values <10 pg/mL were arbitrarily assigned a concentration of 5 pg/mL.

### CD4^+^ T cell depletion

Single cell splenocyte suspensions, obtained as described above, were divided into two groups: mock-depleted and CD4-depleted cells. For the CD4-depleted group, splenocytes were depleted of CD4^+^ T cells using the Mouse CD4 (L3T4) MicroBeads (#130-117-043, Miltenyi Biotec, Bergisch Gladbach, Germany), a MACS LD column (#130-042-901, Miltenyi Biotec), and a QuadroMACS^TM^ separator following the manufacturer’s instructions. Briefly, splenocytes (1 x 10^8^ cells/mL in MACS^®^ Separation buffer (#130-091-221, Miltenyi Biotec) were mixed with CD4 (L3T4) MicroBeads (10% v/v) and incubated for 10 min at 4°C. MACS LD columns were placed in the magnetic field of the QuadroMACS^TM^ separator and rinsed with 2 mL of separation buffer. To remove clumps, cell suspensions were passed through 30 µm pre-separation filters and then transferred onto the pre-rinsed LD column, followed by two subsequent column washes with 1 mL of separation buffer. The unbound (CD4-depleted) cells were collected in the flow-through solution. The mock-depleted group was submitted to the same conditions except only buffer was used with no beads added. Splenocytes in the two groups of cells were then left unstimulated or stimulated with the respective recombinant protein (5 µg/mL) or peptide (2.5 µg/mL). PMA (phorbol 12-myristate 13-acetate) plus ionomycin (IO) at 10 ng/mL and 1 µg/mL, respectively, served as a positive control. Cells were cultured for three days at 37°C with 5% CO2, stored at −80°C, and IFNγ levels in the culture supernatant were measured by ELISA.

### Flow Cytometry

Bovine serum albumin (BSA) was added to 1X PBS at a concentration of 0.5% as FACS buffer for flow cytometry staining. Splenocytes from total (sham-depleted) or CD4-depleted cells, at 1 x 10^6^ cells per sample, were washed with FACS buffer and stained for 30 min with LIVE/DEAD Fixable Green dead cell stain kit (#L34970, Invitrogen by Thermo Fisher Scientific) according to the company’s instructions. Next, cells were washed and subsequently stained for 30 min in the dark at 4°C with the anti-mouse cell surface markers CD45-APC/Cyanine7 (Clone: QA17A26), CD3-PE (Clone: 145-2C11), CD4-PerCP/Cyanine5.5 (Clone: GK1.5), and CD8-APC (Clone: 53-6.7). All conjugated antibodies were from BioLegend (San Diego, CA) and used for staining at a dilution of 1:200. After staining, cells were washed, fixed with 2% paraformaldehyde (PFA), and acquired with a 5-laser Bio-Rad ZE5 Cell Analyzer flow cytometer (Bio-Rad). Data analysis was performed with FlowJo Software, version 10.8 (BD, Franklin Lakes, NJ) and gating was established using FMO controls.

### Data analysis

GraphPad Prism, version 10.1.2 (GraphPad Software, La Jolla, CA) was used for graphing the data and statistical analysis. The IEDB website (www.IEDB.org) was the source for tools to predict IC50 values for peptides sequentially spanning the Cda2 and Cda1 protein sequences. NetMHCIIpan 4.1 BA was used to assign IC50 values to each 15 amino acid peptide and deduce 9 amino acid core sequences for MHC-II epitopes. One-way analysis of variants (ANOVA) was used to compare splenocyte IFNγ responses following stimulation with 15 aa peptides. Kaplan-Meier survival curves were compared using the Mantel-Cox log-rank test; P-values were based on pair-wise comparisons.

### Epitope information

Epitope information has been uploaded to the IEDB (www.iedb.org). The Cda2 Reference IDs in IEDB are: 1040799, 1040334, 1047809, and 1048034. The Cda1 Reference IDs in IEDB are: 1040804, 1041101, and 1047550.

## Acknowledgments

The authors thank Drs. Jane Homan, ChiungYu Hung, Althea Campuzano, Bruce Klein, Marcel Wuethrich, George Deepe, and Melanie Trombly for helpful suggestions and feedback during our monthly scientific teleconferences. This work was supported by grants R24AI192252 (SML), R01AI172154 (SML), and R01AI125045 (SML, CAS), and contract 75N93019C00064, all awarded by the National Institute of Allergy and Infectious Diseases. The funders had no role in study design, data collection and interpretation, or the decision to submit the work for publication.

CAS helped develop the study, analyze data, and write the manuscript. LVNO helped develop the study, perform assays, analyze data, and write the manuscript. DC helped develop the study, perform assays and analyze data. MMH helped develop the study, perform assays, analyze data, and write the manuscript. SS helped perform assays, analyze data, and write the manuscript. GKP supplied reagents and scientific input. SML helped develop the study, analyze data, and write the manuscript. All authors read and approved the content of the final manuscript. CAS and LVNO contributed equally to this manuscript.

## References

1. Rajasingham R, Govender NP, Jordan A, Loyse A, Shroufi A, Denning DW, Meya DB, Chiller TM, Boulware DR. 2022. The global burden of HIV-associated cryptococcal infection in adults in 2020: a modelling analysis. The Lancet Infectious Diseases 22:1748–1755.

2. Denning DW. 2024. Global incidence and mortality of severe fungal disease. Lancet Infect Dis doi:10.1016/S1473-3099(23)00692-8.

3. Oliveira LVN, Wang R, Specht CA, Levitz SM. 2021. Vaccines for human fungal diseases: close but still a long way to go. NPJ Vaccines 6:33.

4. Del Poeta M, Wormley FL, Jr., Lin X. 2023. Host populations, challenges, and commercialization of cryptococcal vaccines. PLoS Pathog 19:e1011115.

5. Avina SL, Pawar S, Rivera A, Xue C. 2024. Will the Real Immunogens Please Stand Up: Exploiting the Immunogenic Potential of Cryptococcal Cell Antigens in Fungal Vaccine Development. Journal of Fungi 10:840.

6. Oliveira LVN, Hargarten JC, Wang R, Carlson D, Park Y-D, Specht CA, Williamson PR, Levitz SM. 2025. Peripheral blood CD4+ and CD8+ T cell responses to Cryptococcus candidate vaccine antigens in human subjects with and without cryptococcosis. Journal of Infection 91:106521.

7. Carlson D, Wang R, Hastings Z, Oliveira LVN, Hester MM, Rodriguez N, Pedersen GK, Tenor JL, Perfect JR, Specht CA, Levitz SM. 2025. Vaccine-Mediated Protection of Mice Against African and Asian Clinical Strains of Cryptococcus neoformans. J Fungi (Basel) 11.

8. Wang R, Oliveira LVN, Hester MM, Carlson D, Christensen D, Specht CA, Levitz SM. 2024. Protection against experimental cryptococcosis elicited by Cationic Adjuvant Formulation 01-adjuvanted subunit vaccines. PLOS Pathogens 20:e1012220.

9. Wang R, Oliveira LVN, Lourenco D, Gomez CL, Lee CK, Hester MM, Mou Z, Ostroff GR, Specht CA, Levitz SM. 2023. Immunological correlates of protection following vaccination with glucan particles containing Cryptococcus neoformans chitin deacetylases. NPJ Vaccines 8:6.

10. Hester MM, Lee CK, Abraham A, Khoshkenar P, Ostroff GR, Levitz SM, Specht CA. 2020. Protection of mice against experimental cryptococcosis using glucan particle-based vaccines containing novel recombinant antigens. Vaccine 38:620–626.

11. Specht CA, Lee CK, Huang H, Hester MM, Liu J, Luckie BA, Torres Santana MA, Mirza Z, Khoshkenar P, Abraham A, Shen ZT, Lodge JK, Akalin A, Homan J, Ostroff GR, Levitz SM. 2017. Vaccination with Recombinant Cryptococcus Proteins in Glucan Particles Protects Mice against Cryptococcosis in a Manner Dependent upon Mouse Strain and Cryptococcal Species. MBio 8:e01872–17.

12. Specht CA, Wang R, Oliveira LVN, Hester MM, Gomez C, Mou Z, Carlson D, Lee CK, Hole CR, Lam WC, Upadhya R, Lodge JK, Levitz SM. 2024. Immunological correlates of protection mediated by a whole organism, Cryptococcus neoformans, vaccine deficient in chitosan. mBio doi:10.1128/mbio.01746-24:e0174624.

13. Pishesha N, Harmand TJ, Ploegh HL. 2022. A guide to antigen processing and presentation. Nature Reviews Immunology 22:751–764.

14. Leddy O, Yuki Y, Carrington M, Bryson BD, White FM. 2025. Targeting infection-specific peptides in immunopeptidomics studies for vaccine target discovery. J Exp Med 222.

15. Sofron A, Ritz D, Neri D, Fugmann T. 2016. High-resolution analysis of the murine MHC class II immunopeptidome. European Journal of Immunology 46:319–328.

16. Racle J, Guillaume P, Schmidt J, Michaux J, Larabi A, Lau K, Perez MAS, Croce G, Genolet R, Coukos G, Zoete V, Pojer F, Bassani-Sternberg M, Harari A, Gfeller D. 2023. Machine learning predictions of MHC-II specificities reveal alternative binding mode of class II epitopes. Immunity 56:1359–1375.e13.

17. Homan EJ, Bremel RD. 2023. Determinants of tumor immune evasion: the role of T cell exposed motif frequency and mutant amino acid exposure. Frontiers in Immunology 14.

18. Sant’Angelo DB, Robinson E, Janeway CAJ, Denzin LK. 2002. Recognition of core and flanking amino acids of MHC class II-bound peptides by the T cell receptor. European Journal of Immunology 32:2510–2520.

19. Arrieta-Bolanos E, Hernandez-Zaragoza DI, Barquera R. 2023. An HLA map of the world: A comparison of HLA frequencies in 200 worldwide populations reveals diverse patterns for class I and class II. Front Genet 14:866407.

20. Gonzalez-Galarza Faviel F, McCabe A, Santos Eduardo J Md, Jones J, Takeshita L, Ortega-Rivera Nestor D, Cid-Pavon Glenda MD, Ramsbottom K, Ghattaoraya G, Alfirevic A, Middleton D, Jones Andrew R. 2019. Allele frequency net database (AFND) 2020 update: gold-standard data classification, open access genotype data and new query tools. Nucleic Acids Research 48:D783–D788.

21. Peters B, Nielsen M, Sette A. 2020. T Cell Epitope Predictions. Annual Review of Immunology 38:123–145.

22. Vita R, Blazeska N, Marrama D, Members ICT, Duesing S, Bennett J, Greenbaum J, De Almeida Mendes M, Mahita J, Wheeler Daniel K, Cantrell Jason R, Overton James A, Natale Darren A, Sette A, Peters B. 2024. The Immune Epitope Database (IEDB): 2024 update. Nucleic Acids Research 53:D436–D443.

23. Sahu SR, Specht CA, Levitz SM. 2026. How post-translational modifications in pathogenic fungi inform pathogenesis and immune responses. PLoS Pathog 22:e1014162.

24. Ito K, Bian HJ, Molina M, Han J, Magram J, Saar E, Belunis C, Bolin DR, Arceo R, Campbell R, Falcioni F, Vidović D, Hammer J, Nagy ZA. 1996. HLA-DR4-IE chimeric class II transgenic, murine class II-deficient mice are susceptible to experimental allergic encephalomyelitis. The Journal of experimental medicine 183:2635–2644.

25. Specht CA, Homan JE, Lee CK, Mou Z, Gomez CL, Hester MM, Abraham A, Rus F, Ostroff GR, Levitz SM. 2021. Protection of mice against experimental cryptococcosis by synthesized peptides delivered in glucan particles. mBio 13:e033674.

26. Pedersen GK, Andersen P, Christensen D. 2018. Immunocorrelates of CAF family adjuvants. Semin Immunol 39:4–13.

27. Wiesner DL, Specht CA, Lee CK, Smith KD, Mukaremera L, Lee ST, Lee CG, Elias JA, Nielsen JN, Boulware DR, Bohjanen PR, Jenkins MK, Levitz SM, Nielsen K. 2015. Chitin recognition via chitotriosidase promotes pathologic type-2 helper T cell responses to cryptococcal infection. PLoS pathogens 11:e1004701.

28. Hain S, Fu MS, Wigg L, George L, Lecky D, Whitehead AJ, Clipston E, Sato K, Ono M, Wuthrich M, Klein B, Kawakami K, Rayes J, Bending D, Drummond RA. 2025. Brain-infiltrating CD4 T cells drive inflammatory microglia proliferation during cryptococcal meningitis in mice. Nature Communications 16:8995.

29. Peters B, Nielsen M, Sette A. 2020. T Cell Epitope Predictions. Annual Review of Immunology 38:123–145.

30. Paul S, Grifoni A, Peters B, Sette A. 2020. Major Histocompatibility Complex Binding, Eluted Ligands, and Immunogenicity: Benchmark Testing and Predictions. Frontiers in Immunology Volume 10 - 2019.

31. Malandro N, Budhu S, Kuhn Nicholas F, Liu C, Murphy Judith T, Cortez C, Zhong H, Yang X, Rizzuto G, Altan-Bonnet G, Merghoub T, Wolchok Jedd D. 2016. Clonal Abundance of Tumor-Specific CD4+ T Cells Potentiates Efficacy and Alters Susceptibility to Exhaustion. Immunity 44:179–193.

32. Liu J, Wei H, Liu J, Peng L, Li G, Li M, Yang L, Jiang Y, Peng F. 2022. Analysis of the Association of HLA Subtypes With Cryptococcal Meningitis in HIV-Negative Immunocompetent Patients. Future Microbiology 17:1231–1240.

33. Skipper CP, Dai B, Borges B, Augusto DG, Gerlach E, Tukundane A, Tadeo K, Pastick KA, Williams DA, Kabahubya M, Rajasingham R, Nalintya E, Meya DB, Boulware DR, Hollenbach J, Nielsen K. 2026. Investigation of human leukocyte antigen alleles as risk factors for cryptococcal disease in Ugandan individuals with HIV. Human Immunology 87:111788.

34. Chen X, Zaro JL, Shen W-C. 2013. Fusion protein linkers: Property, design and functionality. Advanced Drug Delivery Reviews 65:1357–1369.

35. Hurtgen BJ, Hung CY, Ostroff GR, Levitz SM, Cole GT. 2012. Construction and evaluation of a novel recombinant T cell epitope-based vaccine against Coccidioidomycosis. Infect Immun 80:3960–74.

36. Coelho MA, David-Palma M, Aylward J, Pham NQ, Visagie CM, Fuchs T, Yilmaz N, Roets F, Sun S, Taylor JW, Wingfield BD, Fisher MC, Wingfield MJ, Heitman J. 2025. Decoding Cryptococcus: From African biodiversity to worldwide prevalence. PLOS Pathogens 21:e1012876.

37. Nielsen K, Cox GM, Wang P, Toffaletti DL, Perfect JR, Heitman J. 2003. Sexual cycle of Cryptococcus neoformans var. grubii and virulence of congenic a and alpha isolates. Infection and immunity 71:4831–41.

